# Temporal variation and drivers of *Ixodiphagus hookeri* parasitisation in *Ixodes ricinus* ticks in northern Europe

**DOI:** 10.64898/2026.01.14.699597

**Authors:** Jani J. Sormunen, Laura Väisänen, Jesse Mänttäri, Ilari E. Sääksjärvi, Eero J. Vesterinen, Tero Klemola

## Abstract

*Ixodiphagus hookeri* (Hymenoptera: Encyrtidae) is a highly specialized koinobiont endoparasitoid of hard ticks (Acari: Ixodidae). While it is considered a prime target for biological control of ticks, longitudinal data on natural parasitisation of tick populations is scarce, consequently leading to a lack of understanding regarding the effects of global warming on wasp populations and factors impacting natural parasitisation rates of ticks. To fill in these gaps, we present a longitudinal time series on natural *I. hookeri* parasitisation of *Ixodes ricinus* from the northernmost limits of their occurrence.

In total, 2,111 *I. ricinus* nymphs collected between 2014-2021 from Seili Island in southwestern Finland were screened for *I. hookeri* DNA utilizing an in-house qPCR assay. Samples had previously been screened for several tick-borne pathogens and larval blood meal sources. Generalized linear mixed models were fitted to determine factors influencing the probability of parasitisation of questing nymphs, whether parasitisation influences tick densities, and if differences in parasitisation can be identified between early (May to mid-July) and late (mid-July to October) season. Log odds ratios were calculated to assess parasitisation-pathogen and parasitisation-host associations. An artificial tick feeding system was used to feed nymphs collected from the island.

An increase in *I. hookeri* parasitisation was observed on the island between 2014-2021. The overall parasitisation prevalence was 8.2 %, with year and study site specific values ranging from 0 to 38 %. Larvae densities, July mean temperatures and parasitisation rates at *t*-1 were observed to increase probability of parasitisation. Parasitisation was more likely if the nymphs had fed on deer as larvae or carried *Babesia* spp. protozoa, and less likely if they carried *Borrelia* spp. Twelve wasps emerged from an artificially fed nymph, immediately starting oviposition and expiring within seven days.

We present here the longest time series of natural *I. hookeri* parasitisation of *I. ricinus* ticks. An increasing trend in parasitisation was observed, indicating increases in the wasp population likely linked to global warming. Our results regarding associations between parasitisation and tick-borne pathogens - and directly linking parasitisation events to deer - support findings from previous research, which have indicated that parasitisation of ticks is more likely to occur on large host animals than on small hosts or ticks questing on the ground. The results suggest that larvae may be more important parasitisation targets for the wasps than previously accredited.

## Introduction

*Ixodiphagus hookeri* is a parasitoid wasp (Hymenoptera: Encyrtidae) specialized in parasitising hard ticks (Acari: Ixodidae). It has been reported to parasitise 21 different species of ticks and to have a nearly cosmopolitan distribution, being detected from, *e*.*g*., North and South America, Europe, Africa, Southeast Asia, India, and Australia [1, 2]. The wasps are gregarious koinobiont endoparasitoids and parasitise tick larvae and nymphs. The development of wasp eggs commences during the nymphal blood meal, but not during the larval blood meal [3, 4]. Successfully parasitised engorged nymphs die prior to becoming adults and reproducing, which has led to interest towards the wasp as a biological control agent against ticks [1]. So far, prior field trials testing the utility of the wasp in biological control have not led to groundbreaking outcomes or the wide-scale adoption of the wasp as a control method. However, understanding of *Ixodiphagus* biology has advanced in the years following these field trials, and the increasing abundance of ticks over the past few decades has sparked novel interest in the subject [5-9]. As knowledge regarding the biology and phenology of these parasitoids increases, potential shortcomings in previous field trials may also have been uncovered.

Field trials regarding the control potential of *I. hookeri* have been conducted in Kenya and the USA, with mixed success. In field trials in Kenya, results were promising, but highlighted that not all tick species are parasitised by local wasps [10]. Supporting this finding, laboratory studies have suggested that European *I. hookeri* readily parasitise *I. ricinus*, but not another locally occurring tick species, *Dermacentor reticulatus* [3]. Indeed, while wasps parasitising various tick species across the globe are synonymously called *I. hookeri*, it is likely that they are in fact different species or at least subspecies [3]. It is possible that specific wasp species/subspecies are specialized in parasitising only specific tick species, consequently being less effective in parasitising others - or even unable to. In addition, the wasps appear to also show local adaptations in, *e*.*g*., phenologies, as exemplified by the fact that adult wasps were observed to have only short flight periods in the temperate climate of Germany, whereas they were observed to be active throughout the year in tropical Kenya [3, 4]. Any of these factors – incorrect wasp species/subspecies, host specificity towards non-native tick species, mismatching local adaptations - may have influenced results obtained in the USA, which have been less promising than those obtained from Kenya [11-14]. At least on one occasion wasps sourced from France were used in field trials, whose ability to parasitise the local tick species may have been limited [14]. It is also unclear what sort of an effect releases that are not matched to the varying natural phenology of the wasps might have on the success of control efforts. Indeed, there is very scarce data on the phenology of *I. hookeri* in general, hindering efforts to utilize them for control [3, 4]. Locally sourced wasp populations utilized in concordance with their phenology are the most likely to lead to successful control actions, highlighting the need for localized data.

Laboratory studies have indicated that odours from tick host animals – rather than odours from ticks themselves – act as the major attractants for *I. hookeri* [9, 15, 16]. Specifically, *I. hookeri* parasitising *I. ricinus* have been observed to be attracted towards deer and wild boar odours, but not those of cattle, rabbits or mice [15]. This indicates that the wasps seek out large wildlife hosts in order to find ticks, rather than searching for questing ticks on the ground [17]. *Ixodiphagus hookeri* are tiny chalcidoid wasps with short life spans [17], typically moving by jumping short distances rather than by flying. This renders searching for ticks to parasitise on the ground ineffective for maximizing reproductive potential. In contrast, scores of ticks can be found aggregated on their hosts. Likewise, ticks on hosts are more likely to survive than questing ticks, since they have already found a blood meal source [15]. A recent study from the Netherlands highlights the importance of large wildlife hosts as parasitisation hubs, observing a positive association between deer densities and parasitisation rates [18]. Furthermore, a study from Connecticut, USA, observed that *I. hookeri* parasitisation of *I. scapularis* was reduced following a reduction in white-tailed deer densities [19]. Finally, a study on *I. hookeri* parasitisation of *Amblyomma variegatum* in Kenya showed that only ticks collected from cattle were parasitised, but not those collected from the grass [17]. These studies indicate that parasitisation may happen mostly on host animals. However, if ticks are mostly parasitised on hosts, larvae may in fact be major parasitisation targets, even though laboratory studies have indicated that wasps prefer nymphs [3, 18]. This is because the parasitisation of nymphs already on hosts is not likely to lead to the commonly detected parasitised questing nymphs [18, 20].

The northernmost observations of *I. hookeri* have been made from Finland and Norway. In Finland, adult wasps have been reported on two occasions, in the 1950’s from Lohja by Wolter Hellén [21] and in 2013 from Seili Island [20], both located at the southernmost reaches of the country. While only one adult wasp was detected from Seili by Malaise trapping in 2013, molecular analysis of ticks collected from the island between 2012-2014 revealed parasitisation of questing *I. ricinus* nymphs therein [20]. In 2017, an effort to estimate the flight periods of adult wasps was made on Seili Island, utilizing sweep netting and Malaise traps [20]. However, no additional adult wasps were detected. As such, the closest estimate regarding the flight periods of adult wasps is from Germany, where adult *I. hookeri* were observed to be active for 3-5 weeks in mostly July and August [3]. In the study from Germany, cloth dragging rather than sweep netting was used to sample wasps [3], which may be a more suitable method. However, the recent detection of an adult wasp from Norway was made by sweep netting [2]. Both of the more recent detections of adult wasps from Finland and Norway were made between late June and early August, which implies that *I. hookeri* activity is likely to follow a similar trend as in Germany. In contrast to observations from Europe, adult wasps were observed throughout the year in Kenya [4], indicating that lower temperatures and the clear seasonality in climate in Europe are limiting wasp activity. Furthermore, the fact that detections from Finland and Norway have only been made from the southernmost parts of the countries implies that temperature may be the main limiting factor for wasp occurrence. Indeed, ticks are found in abundance also further north than these collection locations [22, 23], so occurrence is not limited by the absence of ticks. The effect of temperature on *I. hookeri* occurrence solicits further research, but modelling efforts are currently limited by scarce occurrence data for the wasps.

A previous study conducted on Seili Island in southwestern Finland indicated that *I. hookeri* may have emerged on the study site relatively recently [20]. However, due to a lack of longitudinal data, the development of the local population could not be assessed then. In fact, there is a general scarcity of studies observing natural *I. hookeri* parasitisation in tick populations across time [24], hindering efforts to estimate how wasp populations may be developing and/or fluctuating. This is vital data to uncover the driving forces for changing abundance and distributions, and to assess potential future impacts of the warming climate on wasp populations.

In the current study, we present longitudinal data from 2014-2021 regarding *I. hookeri* parasitisation of questing *I. ricinus* nymphs at Seili Island in southwestern Finland. We assess how temperature, rainfall, tick densities and parasitisation observed in the previous year influence parasitisation probability. Furthermore, we utilize previous data on tick-borne pathogens and larval blood meal sources of the analysed nymphs to assess parasitisation-pathogen and parasitisation-host relationships. Finally, we describe the emergence of *I. hookeri* from an artificially fed nymph collected from the study island and share observations made regarding their biology.

## Materials and methods

### Tick samples and pathogen analysis

Tick samples collected from an island in southwestern Finland, Seili (60.23° N, 21.96° E), between 2014-2021 were utilized in this study [25]. Tick studies have been conducted on this rural island since 2012 and the first national report of *I. hookeri* parasitisation of *I. ricinus* was also made therein [20, 25]. Tick collection and pathogen screening methods have been described in detail in previous work [25, 26]. Briefly, study sites on the island take the form of 50 meter fixed transects, which have been dragged 1-3 times monthly mainly from May to September. For this study, one transect in deciduous forest, two transects in coniferous forest and one transect in an alder thicket were chosen. Recent efforts in analysis of samples from the island have focused on these study lines, which tend to have the highest densities of ticks and thus supply sufficient ticks for laboratory analyses [27]. Consequently, extracted DNA was available mostly for these study sites.

DNA and RNA have been extracted from the ticks utilizing Macherey-Nagel RNA kits with DNA buffer sets (Macherey-Nagel, Germany) [26]. The majority of the tick samples utilized in this study were previously screened for tick-borne pathogens *Borrelia* spp., *Rickettsia* spp., *Babesia* spp., *Neoehrlichia mikurensis, Anaplasma phagocytophilum, Bartonella* spp. and *Francisella tularensis* using multiplex qPCR assays [26].

### Molecular detection of Ixodiphagus hookeri and blood meal analysis

Tick parasitisation on the island was previously studied for the years 2012-2014 utilizing qPCR and Sanger sequencing [20]. In this previous research, samples from 2012-2013 were pooled, so only minimum infection rates were available for these years. However, closer inspection of results from 2014 revealed that all the nymphs from the chosen study sites were analysed separately, so 2014 data was incorporated into analyses. For samples from 2015-2021, *I. hookeri* parasitisation was analysed using an in-house qPCR assay as previously described [20]. DNA extracts from *I. hookeri* that emerged from the laboratory-fed *I. ricinus* nymph were used as positive controls.

Tick samples collected from Seili in 2021 were previously analysed for nymph blood meal sources utilizing retrotransposon-based qPCR methods [27-30]. However, since fewer ticks had been analysed for deer than other host groups and because we observed a potential positive association between parasitisation and deer as hosts, we analysed an additional 50 *I. hookeri* positive and negative samples collected in 2018-2021 with the deer blood meal assay.

### Artificial tick feeding

We used a previously published artificial tick feeding system utilizing silicone membranes to feed nymphs collected from the island [31, 32]. We utilized membranes of 50-70 µm thickness and a fat extract created from the hair of white-tailed deer (*Odocoileus virginianus*). A total of 150 nymphs collected from Seili were divided across six feeding units, 25 nymphs per unit. Out of these, 68 successfully fed to repletion before feeding was stopped.

### Statistical analyses

The main objective for statistical analyses was to identify factors influencing parasitisation probability of questing nymphs. The flight period of adult *I. hookeri* is expected to be very short locally, so we hypothesized that weather conditions during this short flight period could have a major impact on the success of parasitisation each year. While the exact flight times in Finland remain unclear, they are likely to be in July and August, as observed in Germany [3]. As such, we assessed the effects of temperature and rainfall for not only the main tick activity season (May to September) but also July and August separately (including their means). Regarding tick densities, larvae and nymphs are the targets of the wasps, so only larvae and nymph densities were included in analyses. It is possible that tick densities during the flight period in the late season (July-September) may be more important for parasitisation, so we included study site specific late season (week 29 onwards) densities in addition to total activity season densities, separately for each life stage. Furthermore, we included a larvae-nymph density ratio (larvae/nymphs) for both the main activity period and late season, in case the parasitisation of larvae is dependent on their relative availability compared to nymphs. Finally, we included parasitisation rates of the previous year (*t*-1), again separately for each study site.

We coded the binary parasitisation data as binomial successes-trials data. Consequently, we were left with 32 unique observation groups (4 study sites, 8 years). In order to keep our ratio of observations to explanatory variables as close to the preferred ≥10:1 as possible [33], we limited our model to a maximum of four explanatory variables. For model selection, we z-standardized all the variables and divided them into four groups: temperature data, rainfall data, tick density data and parasitisation data, all at *t*-1. We then used an R script to run generalized linear mixed models (GLMMs) containing each possible combination of explanatory variables (*n*=350; also including the possibility to leave the group out from the model) and a random effect for study site, and selected the model that was most supported by AIC and R^2^ (Table S1). The best performing model included larvae density, July mean temperature and parasitisation at *t-*1, as well as the study site random effect and the logit link function.

To determine if parasitisation during the previous year (*t*-1) had an effect on tick densities at *t*, we ran generalized linear mixed models where densities of specific tick life stages (larvae, nymph, and adult) from 2015 to 2021 were explained by parasitisation rates at *t*-1. Each transect on the island is exactly 50 meters and are dragged the same number of times, so no offset variables were required for the model. Only minimum infection rate data for previous parasitisation rates was available for 2014, so the year was dropped from the analysis. Finally, we utilized a binomial GLMM, with the logit link function, to analyse differences in parasitisation probabilities of nymphs collected in the early (weeks 19-28) or late (29-42) season. Study site was used as a random effect in all the models. Log odds ratios were calculated for binary parasitisation–host and parasitisation–pathogen interactions to assess associations.

Generalized linear mixed models were run using R Statistical Software (v4.5.2) [34]. We utilized packages *lme4* [35], *purrr* [36] and *dplyr* [36] for GLMMs and analysis, and package *performance* [37] to calculate R^2^ for GLMMs. Log odds ratios were calculated using the JASP software (version 0.17.2.1; https://jasp-stats.org/). All figures were created using package *ggplot2* [38] in RStudio.

## Results

In total 174 *I. ricinus* nymphs parasitised by *I. hookeri* were detected among 2,111 analysed samples from 2014 to 2021, with an overall parasitisation prevalence of 8.2 ± 1.2 % (with 95% confidence limits). An increasing trend in parasitisation was observed, although interannual variation was high (Figure 1). Study site and year specific parasitisation prevalence ranged from 0 to 38 % (Table S2). In addition to the overall increase in parasitisation, parasitised nymphs were observed on two of the four study transects only from 2017 or 2018 onwards (Table S2), indicating also distributional changes within the island.

**Figure 1.**
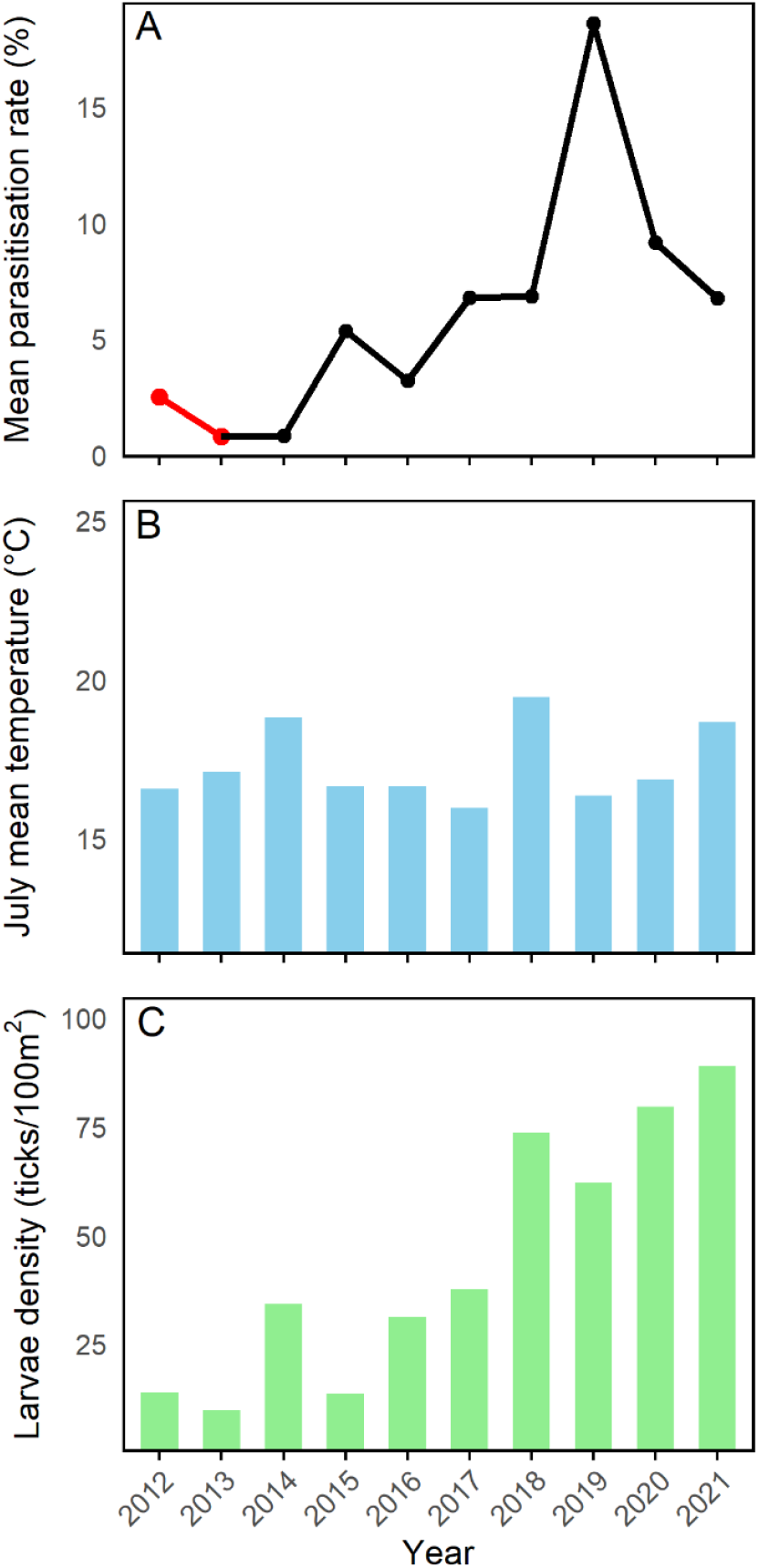
Annual mean Ixodiphagus hookeri parasitisation rates in questing Ixodes ricinus nymphs (A), July mean temperature (B), and I. ricinus larvae density (C) on Seili Island, 2012-2021. Please note that parasitisation rates for 2012 and 2013 are minimum infection rates (highlighted in red).

Larvae densities, July mean temperatures, and parasitisation rates during the previous year (*t-*1) were all observed to have a positive effect on parasitisation probability (Table 1, Figure 2). Out of the 350 models ran, overall larvae densities were included in the five best performing models, July mean temperature in nine, and previous parasitisation rates in one hundred and five. However, a temperature variable was included in two hundred and thirty-eight of the best performing models, highlighting the relevance of temperature (Table S1). The best performing model obtained marginal and conditional R^2^ values of 0.85 and 0.92, indicating that the fixed factors in the model explained most of the observed variation in parasitisation probability, with a minor contribution from unassigned spatial variation.

**Table 1.** Results from the binomial model estimating the effects of I. ricinus larvae density, July mean temperature and parasitisation by Ixodiphagus hookeri in the previous year on the probability of parasitisation for questing nymphs.

| Variable | Estimate | 95% CI<sup>a</sup> | z-value | p |
| --- | --- | --- | --- | --- |
| Intercept | -2.8 | -3.1 to -2.5 | -18.5 | <0.0001 |
| Larvae density<sub>t-1</sub> | 0.25 | 0.05 to 0.45 | 2.4 | 0.01 |
| July temperature<sub>t-1</sub> | 0.62 | 0.46 to 0.78 | 7.9 | <0.0001 |
| Parasitisation<sub>t-1</sub> | 0.50 | 0.30 to 0.70 | 5.2 | <0.0001 |
|  | Variance | Standard deviation |  |  |
| Study Site (random effect) | 0.04 | 0.21 |  |  |

<sup>a</sup>95% confidence limits

**Figure 2.**
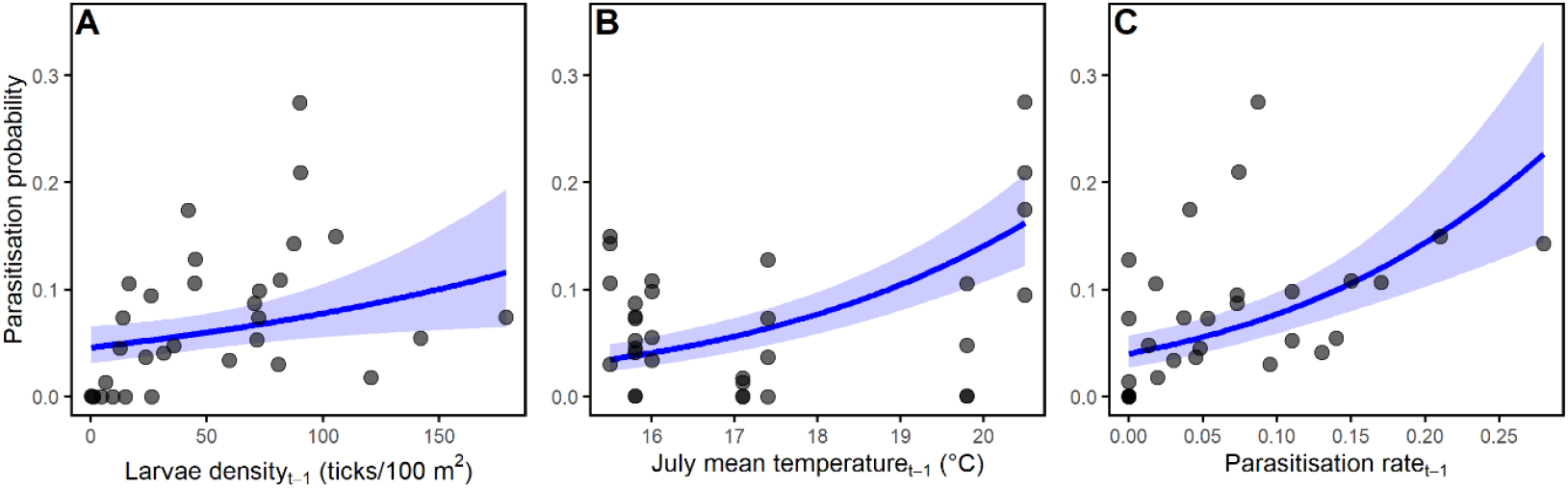
Observed Ixodiphagus hookeri parasitisation prevalence in questing Ixodes ricinus nymphs (dots) and predictions of parasitisation probability from the statistical model (lines with 95% confidence limits). Presented for I. ricinus larvae density (A), July mean temperature (B), and parasitisation rates (C), all at year t-1.

Parasitisation probabilities of the previous year were positively association with larvae densities (β = 0.48 ± 0.21 SE, *z* = 2.3, *p* = 0.02) and not associated with other life stages. Likewise, no differences were observed between early [0.054 (0.025 – 0.16)] and late [0.056 (0.026 – 0.16)] season parasitisation probability for questing nymphs (β = 0.04 ± 0.18 SE, *z* = 0.21, *p* = 0.84).

Log odds ratios revealed a higher chance to be parasitised if the nymph had previously fed on deer or carried *Babesia* spp. protozoa (Figure 3). Contrastingly, carrying *Borrelia* spp. bacteria was observed to lower the chance to be parasitised (Figure 3).

**Figure 3.**
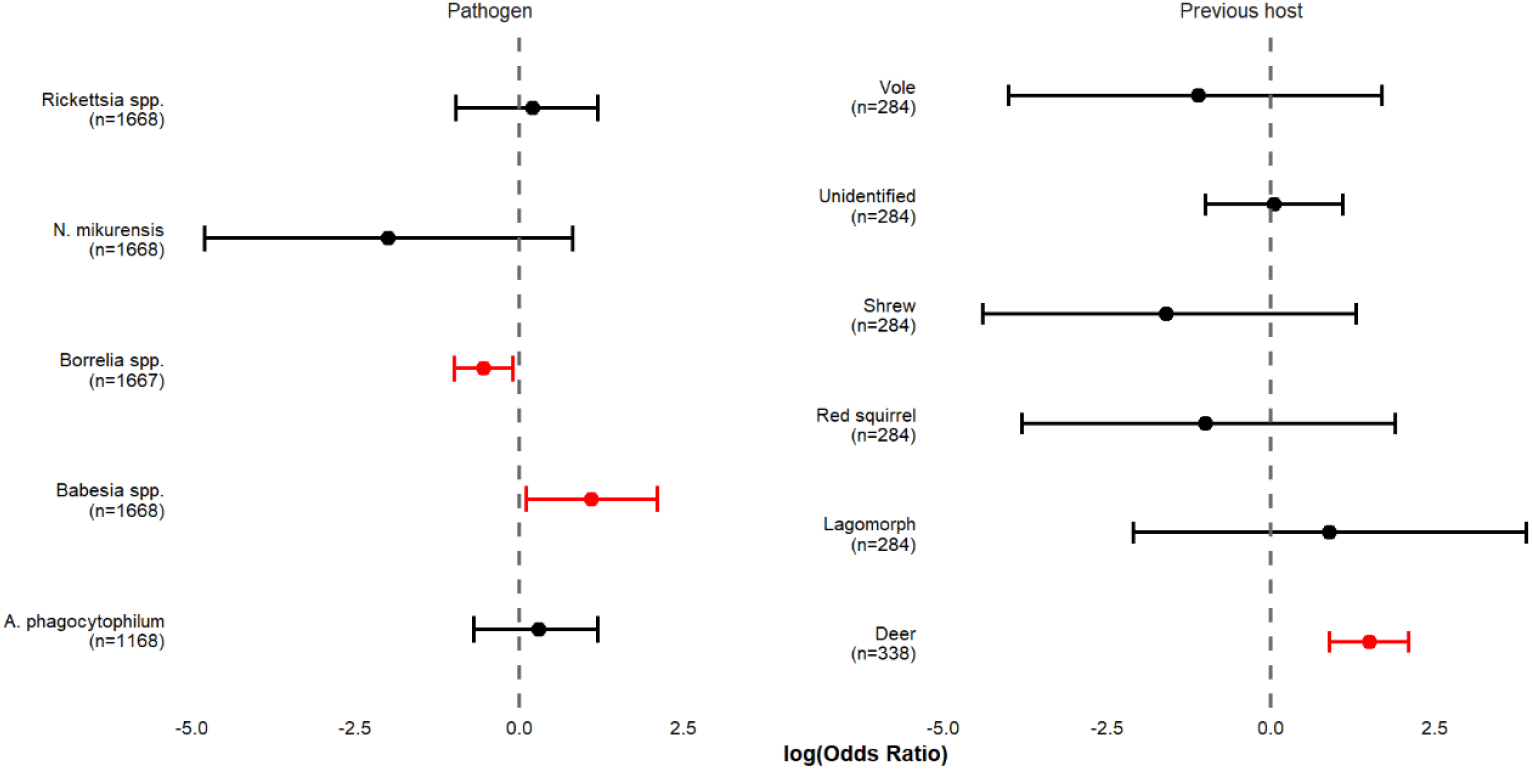
Log odds ratios for associations of Ixodiphagus hookeri parasitisation with tick-borne pathogens carried by and source of the larval blood meal of questing Ixodes ricinus nymphs. Number of samples available for analyses is reported in brackets for each pair. Significant odds ratios are highlighted in red.

Twelve *Ixodiphagus hookeri* emerged from an *I. ricinus* nymph collected from Seili island that was fed to repletion using an artificial tick feeding system. The tick had finished feeding on 11 June 2024 and was held in darkness at above 95 % relative humidity and room temperature (21-23 ℃). The wasps emerged 35 days later, on 16 July 2024. During feeding, the tick was observed to become larger than non-parasitized nymphs, but no weighing was conducted. During development, the tick was observed to become brownish, until a routine examination of developing nymphs on 16 July 2024 revealed a powder like substance on the bottom of the sample tube. Microscope inspection revealed a wasp gnawing a hole in the posterior end of the tick as the source of the substance. Roughly 30 minutes later, all the wasps had emerged from the tick. The wasps were placed in a sample tube with roughly 20 *I. ricinus* nymphs within a few hours of their emergence and immediately started to oviposition once they encountered the ticks. The wasps were maintained at room temperature and above 95 % relative humidity (tick storage conditions) with the ticks. The last wasp died on 22 July 2024, indicating a life span of seven days at maximum for adult wasps. However, it is unknown how the antibiotic (gentamicin) fed to the ticks during feeding might influence wasp longevity. Unfortunately, subsequent feeding of parasitised nymphs failed due to a fungal contamination in the feeding unit, which led to the loss of the entire group.

## Discussion

The current study revealed a longitudinal increase in *I. hookeri* parasitisation of *I. ricinus* nymphs at what is expected to be their northernmost limits of occurrence in Europe. At the same time, the spatial distribution of the wasps seems to have expanded within the study island. These changes in the wasp population seem to be at least partly driven by higher summer temperatures, indicating global warming as a potential cause for changes. Global warming has already been observed to have caused changes in environmental conditions sufficient to impact tick biology in Europe, as evidenced by the first detections of adult *Hyalomma marginatum* in Sweden and the UK in 2018, following a particularly warm summer [39, 40]. Indeed, the highest *I. hookeri* parasitisation rates on Seili were observed in 2019, following the warm summer of 2018, but rates fell again in 2020 and 2021. This might indicate that the growth of the wasp population at these northern latitudes is indeed limited by temperature. Consequently, a warming climate may enable *I. hookeri* to reach higher population densities locally and to expand their distribution range further north. So far, the only reported detections of the wasp from the Nordic countries have been from the southernmost parts of Finland and Norway [2, 20].

Apart from July temperatures, the density of larvae and parasitisation rates in the previous year were observed to have a positive effect on the probability of parasitisation. The effect of previous parasitisation seems relatively obvious – higher parasitisation rates during the previous year lead to higher numbers of adult wasps, which in turn support higher parasitisation rates - even if weather or other conditions during the flight period are suboptimal. However, despite the indication of past parasitisation as a highly significant variable by its inclusion in over a hundred of the best performing models, the falling of parasitisation rates following the 2019 high point seems to suggest that summer temperatures may limit wasp reproduction success even when lots of ticks are being parasitised. It is possible that wasp egg development is regulated by as-yet undetermined temperature thresholds, with development failing if these thresholds are not reached. Regarding the indication of larvae density as a predictor for parasitisation, it may indicate that larvae are in fact an important parasitisation target [18] or just that larvae serve as an indication of overall tick population size and thus the availability of juvenile ticks for parasitisation. While the ratio of larvae to nymphs did not make it into the final model, it is also possible that a positive association with larvae density is indicative of high numbers of larvae leading to more larvae being parasitised, consequently leading to higher numbers of parasitised questing nymphs next year.

Despite parasitisation prevalence reaching a study site specific maximum of 38 %, parasitisation in the previous year was observed to have a positive association with larvae density and no associations with other life stages. This suggests that the wasps are not able to control tick populations at natural occurrence rates. Since mortality is in any case high in the juvenile tick life stages, the additional mortality of nymphs may not have as large an impact as the mortality of adults directly might have [18]. However, this finding does not rule out the possibility of utilizing the wasps as biological control, as the numbers of wasps released in such approaches raise the rates well above natural occurrence rates. However, it does imply that releases have to be made at regular intervals to maintain an effective level of control [10].

The biology and phenology of European *I. hookeri* remain somewhat unclear. Research conducted in Germany showed that adult wasps were detected only during a few weeks in July-August, indicating an extremely short and well synchronized activity period. While efforts to map flight periods of the wasps on Seili Island failed, the only adult wasp caught was from a Malaise trap that was active in July, 2013 [20]. Likewise, an adult wasp was recently collected in Norway between late June – mid July [2]. Furthermore, the wasps that emerged from the laboratory-fed nymph in the current study only lived for a maximum of seven days, although the antibiotics used to feed the ticks and the lack of food for the wasps may have had an impact on this. However, a study on *I. hookeri* from Kenya showed even shorter life spans for wasps (1-3 days), with the availability of nutrition only expanding this by one day [17]. Nevertheless, we observed that the wasps were ready to parasitise ticks very soon after emergence, which also indicates a short but highly active flight period. Together, these data suggest that the flight periods of the wasps are likely to be similar to those observed in Germany [3]. Verification of phenology will be attempted in the future with cloth dragging.

Importantly, our results highlight the importance of deer for the *I. hookeri* life cycle – parasitisation was observed to be more likely when a tick had fed on a deer as a larva, suggesting that parasitisation commonly occurs on these hosts. In general, the roles of deer as blood meal sources for also tick larvae may have been underestimated previously – a recent study from the same study island showed that up to 37% of questing *I. ricinus* nymphs had fed on deer as larvae [27]. Parasitisation was also observed to be more likely when the tick was infected with *Babesia* spp. protozoa. The *Babesia* species detected from Seili island are all associated with large reservoir animals, again indicating large hosts as sites of parasitisation [26]. In contrast, bacterial *Borrelia* spp. infection was observed to negatively influence parasitisation chance. Consequently, this finding suggests that tick parasitisation does not occur as commonly on small hosts, which serve as the reservoirs for *Borrelia*. Overall, these findings mirror those obtained in studies in the Netherlands [18] and the United States [40], although in the latter research the absence of *Borrelia* spirochetes and *Babesia microti* in parasitised *I. scapularis* nymphs was hypothesized to be due to incompatibility between wasp eggs and the pathogens, rather than as indication of host utilization. However, considering the mounting data on the importance of large animals – particularly deer – as targets for the wasps and for their life cycles, it is more likely that this observation is rather due to the wasps not seeking out the reservoir hosts of these pathogens (rodents).

The short flight period of *I. hookeri* in Europe raises some questions regarding their life histories [3]. Specifically, if ticks are only parasitised for a short period in July/August and on hosts, why are we observing equal parasitisation prevalence in questing nymphs throughout the season? Previous laboratory studies have indicated that engorged nymphs are most readily parasitised by wasps, but it is unclear to what degree these results may be influenced by the laboratory conditions [3, 4]. It may be that the wasps readily parasitise ticks when they encounter them, but that their behaviour in the nature may influence where (on which host, which part of the host etc.) they seek the ticks. In any case, it is unclear how the parasitisation of engorged nymphs (or even non-engorged nymphs) on hosts in July/August would lead to parasitised questing nymphs in the spring. It is conceivable that parasitised engorged nymphs could survive nearly a year in the ground, to produce a new cohort of adult wasps in July-August next year, but again, it would not lead to questing parasitised nymphs. Furthermore, eggs laid in questing *Amblyomma variegatum* nymphs were observed to disappear after two days, although this may be a wasp/tick species specific phenomenon [4]. In contrast, parasitisation of (engorged) larvae would lead to larvae that overwinter as engorged larvae or newly moulted nymphs, explaining why parasitised questing nymphs are detected starting from the following spring. The development and hatching of wasp eggs could then be regulated by specific temperature thresholds, which would synchronize emergence of wasps from nymphs that have fed in autumn and spring/early summer to July/August, traditionally the warmest periods of the summer. A temperature threshold for egg development could also explain why parasitisation rates are higher following especially warm summers – more eggs successfully develop in warmer weather, producing more adult wasps. Supporting this view, an inference towards wasps parasitising larvae rather than nymphs was also made in a recent correlative study in the Netherlands [18].

In conclusion, previous studies have suggested that *I. hookeri* utilize host odour to find ticks to parasitise (on the hosts) [9, 15, 16] and that parasitisation commonly occurs on hosts [17]. Likewise, deer have been observed to have a positive effect on tick parasitisation rates in previous studies [18, 19] and were implicated even more directly as sites of parasitisation in the current study. Furthermore, negative associations between parasitisation rates and the occurrence of pathogens associated with small reservoir hosts were observed here and previously [18, 41]. These findings strongly suggest that wasps actively seek large hosts such as deer, subsequently parasitising ticks found on the host. This makes sense from an evolutionary perspective, since they are far more likely to find sufficiently many ticks to parasitise during their short life spans on the hosts than by searching for questing ticks on the ground [3]. The observation of equal numbers of parasitised nymphs throughout the season despite adult wasps only being active for a short period in July/August raises some questions about how the wasp life cycle as observed in Europe could be completed and synchronized if mainly nymphs are the targets of parasitisation. In contrast, parasitisation of larvae in the late season would offer a simple pathway to questing, parasitised nymphs from spring onwards. Consequently, we hypothesize that larvae feeding on large animals such as deer may actually be important targets for European *I. hookeri*, as also suggested in a recent study from the Netherlands [18]. Further studies are needed to determine how readily engorged larvae are parasitised by local *I. hookeri*, whether wasps show preference towards either tick life stage when both are present, and whether temperature thresholds for wasp egg development can be identified. Artificial tick feeding systems offer a good platform for such efforts.

## Acknowledgements

This research was funded by the Jane and Aatos Erkko Foundation, Jenny and Antti Wihuri Foundation and Research Council of Finland (grant number: 360177).

