## Supplementary material for "Temporal variation and drivers of *Ixodiphagus hookeri* parasitisation in *Ixodes ricinus* ticks in northern Europe": Table S1

**TABLE S1**. Results from generalized linear mixed binomial models utilizing different combinations of variables depicting tick densities, rainfall, mean temperature, and parasitisation rates at t-1 to assess the probability of *Ixodes ricinus* nymphs to be parasitised by Ixodiphagus hookeri.

| Group | Abbrevation | Definition |
|---|---|---|
| ALL | 0 | Group left out of model. |
| 1 | Larvae | Overall density of larvae. Separately for each study site and year. |
| 1 | Nymph | Overall density of nymphs. Separately for each study site and year. |
| 1 | Ldens | Density of larvae in the late season (week 29 onwards). Separately for each study site and year. |
| 1 | Ndens | Density of nymphs in the late season (week 29 onwards). Separately for each study site and year. |
| 1 | LN | Larvae to nymph ratio (larvae/nymphs). Separately for each study site and year. |
| 1 | LNlate | Larvae to nymph ratio in the late season. Separately for each study site and year. |
| 2 | SR | Overall seasonal rainfall (May to September) on the study island. |
| 2 | JR | Overall rainfall in July. |
| 2 | AR | Overall rainfall in August. |
| 2 | meanR | Mean of overall rainfall in July and August. |
| 3 | ST | Overall mean seasonal temperature (May to September). |
| 3 | JT | Mean temperature in July. |
| 3 | AT | Mean temperature in August. |
| 3 | meanT | Mean overall temperature in July and August. |
| 4 | Prevpara | Parasitization rate per study site and year. |

| Model_n | Group1 | Group2 | Group3 | Group4 | AIC | BIC | Marginal_R2 | Conditional_R2 |
|---|---|---|---|---|---|---|---|---|
| 1 | Larvae | 0 | JT | Prevpara | 134,90 | 142,23 | 0,85 | 0,92 |
| 2 | Larvae | AR | JT | Prevpara | 136,77 | 145,56 | 0,85 | 0,92 |
| 3 | Larvae | SR | JT | Prevpara | 136,78 | 145,58 | 0,85 | 0,92 |
| 4 | Larvae | meanR | JT | Prevpara | 136,78 | 145,58 | 0,85 | 0,92 |
| 5 | Larvae | JR | JT | Prevpara | 136,86 | 145,65 | 0,85 | 0,92 |
| 6 | Ndens | 0 | JT | Prevpara | 137,92 | 145,25 | 0,82 | 0,91 |
| 7 | LNlate | 0 | JT | Prevpara | 138,00 | 145,33 | 0,83 | 0,91 |
| 8 | Ldens | 0 | JT | Prevpara | 138,01 | 145,34 | 0,81 | 0,91 |
| 9 | 0 | 0 | JT | Prevpara | 138,54 | 144,41 | 0,80 | 0,91 |
| 10 | Larvae | 0 | ST | Prevpara | 138,96 | 146,29 | 0,82 | 0,92 |
| 11 | Larvae | 0 | meanT | Prevpara | 138,96 | 146,29 | 0,82 | 0,92 |
| 12 | LNlate | AR | JT | Prevpara | 139,38 | 148,17 | 0,83 | 0,91 |
| 13 | LNlate | SR | JT | Prevpara | 139,39 | 148,19 | 0,83 | 0,91 |
| 14 | LNlate | meanR | JT | Prevpara | 139,39 | 148,19 | 0,83 | 0,91 |
| 15 | Ndens | AR | JT | Prevpara | 139,54 | 148,33 | 0,82 | 0,91 |
| 16 | LN | 0 | JT | Prevpara | 139,58 | 146,91 | 0,83 | 0,91 |
| 17 | Ndens | JR | JT | Prevpara | 139,69 | 148,49 | 0,81 | 0,92 |
| 18 | Ldens | SR | JT | Prevpara | 139,76 | 148,56 | 0,82 | 0,90 |
| 19 | Ldens | meanR | JT | Prevpara | 139,76 | 148,56 | 0,82 | 0,90 |
| 20 | Ldens | AR | JT | Prevpara | 139,81 | 148,60 | 0,82 | 0,91 |
| 21 | Ndens | SR | JT | Prevpara | 139,81 | 148,60 | 0,82 | 0,91 |
| 22 | Ndens | meanR | JT | Prevpara | 139,81 | 148,60 | 0,82 | 0,91 |
| 23 | LNlate | JR | JT | Prevpara | 139,83 | 148,62 | 0,83 | 0,91 |
| 24 | Ldens | JR | JT | Prevpara | 139,96 | 148,76 | 0,81 | 0,91 |
| 25 | Larvae | JR | ST | Prevpara | 140,02 | 148,82 | 0,82 | 0,92 |
| 26 | Larvae | JR | meanT | Prevpara | 140,02 | 148,82 | 0,82 | 0,92 |
| 27 | 0 | AR | JT | Prevpara | 140,18 | 147,51 | 0,80 | 0,91 |
| 28 | 0 | SR | JT | Prevpara | 140,20 | 147,53 | 0,80 | 0,90 |
| 29 | 0 | meanR | JT | Prevpara | 140,20 | 147,53 | 0,80 | 0,90 |
| 30 | Ndens | 0 | ST | Prevpara | 140,34 | 147,67 | 0,77 | 0,92 |
| 31 | Ndens | 0 | meanT | Prevpara | 140,34 | 147,67 | 0,77 | 0,92 |
| 32 | 0 | JR | JT | Prevpara | 140,44 | 147,76 | 0,80 | 0,91 |
| 33 | Nymphs | 0 | JT | Prevpara | 140,54 | 147,87 | 0,80 | 0,91 |
| 34 | Larvae | SR | ST | Prevpara | 140,54 | 149,34 | 0,83 | 0,92 |
| 35 | Larvae | meanR | ST | Prevpara | 140,54 | 149,34 | 0,83 | 0,92 |
| 36 | Larvae | SR | meanT | Prevpara | 140,54 | 149,34 | 0,83 | 0,92 |
| 37 | Larvae | meanR | meanT | Prevpara | 140,54 | 149,34 | 0,83 | 0,92 |
| 38 | Larvae | AR | ST | Prevpara | 140,57 | 149,36 | 0,82 | 0,92 |
| 39 | Larvae | AR | meanT | Prevpara | 140,57 | 149,36 | 0,82 | 0,92 |
| 40 | LNlate | 0 | ST | Prevpara | 141,28 | 148,61 | 0,79 | 0,91 |
| 41 | LNlate | 0 | meanT | Prevpara | 141,28 | 148,61 | 0,79 | 0,91 |
| 42 | LN | SR | JT | Prevpara | 141,31 | 150,10 | 0,83 | 0,90 |

| | | | | | | | | |
|---|---|---|---|---|---|---|---|---|
| 43 | LN | meanR | JT | Prevpara | 141,31 | 150,10 | 0,83 | 0,90 |
| 44 | Ldens | 0 | ST | Prevpara | 141,32 | 148,64 | 0,78 | 0,91 |
| 45 | Ldens | 0 | meanT | Prevpara | 141,32 | 148,64 | 0,78 | 0,91 |
| 46 | LN | AR | JT | Prevpara | 141,36 | 150,16 | 0,83 | 0,91 |
| 47 | 0 | 0 | ST | Prevpara | 141,53 | 147,39 | 0,77 | 0,91 |
| 48 | 0 | 0 | meanT | Prevpara | 141,53 | 147,39 | 0,77 | 0,91 |
| 49 | LN | JR | JT | Prevpara | 141,53 | 150,33 | 0,83 | 0,91 |
| 50 | Ndens | JR | ST | Prevpara | 141,86 | 150,65 | 0,78 | 0,91 |
| 51 | Ndens | JR | meanT | Prevpara | 141,86 | 150,65 | 0,78 | 0,91 |
| 52 | Ndens | SR | ST | Prevpara | 141,96 | 150,75 | 0,78 | 0,91 |
| 53 | Ndens | meanR | ST | Prevpara | 141,96 | 150,75 | 0,78 | 0,91 |
| 54 | Ndens | SR | meanT | Prevpara | 141,96 | 150,75 | 0,78 | 0,91 |
| 55 | Ndens | meanR | meanT | Prevpara | 141,96 | 150,75 | 0,78 | 0,91 |
| 56 | Nymphs | AR | JT | Prevpara | 142,17 | 150,97 | 0,80 | 0,91 |
| 57 | Nymphs | SR | JT | Prevpara | 142,19 | 150,99 | 0,81 | 0,90 |
| 58 | Nymphs | meanR | JT | Prevpara | 142,19 | 150,99 | 0,81 | 0,90 |
| 59 | Ndens | AR | ST | Prevpara | 142,20 | 150,99 | 0,78 | 0,92 |
| 60 | Ndens | AR | meanT | Prevpara | 142,20 | 150,99 | 0,78 | 0,92 |
| 61 | LNlate | SR | ST | Prevpara | 142,22 | 151,01 | 0,80 | 0,91 |
| 62 | LNlate | meanR | ST | Prevpara | 142,22 | 151,01 | 0,80 | 0,91 |
| 63 | LNlate | SR | meanT | Prevpara | 142,22 | 151,01 | 0,80 | 0,91 |
| 64 | LNlate | meanR | meanT | Prevpara | 142,22 | 151,01 | 0,80 | 0,91 |
| 65 | Nymphs | JR | JT | Prevpara | 142,43 | 151,23 | 0,80 | 0,91 |
| 66 | Ldens | JR | ST | Prevpara | 142,62 | 151,41 | 0,78 | 0,91 |
| 67 | Ldens | JR | meanT | Prevpara | 142,62 | 151,41 | 0,78 | 0,91 |
| 68 | 0 | SR | ST | Prevpara | 142,77 | 150,10 | 0,77 | 0,90 |
| 69 | 0 | meanR | ST | Prevpara | 142,77 | 150,10 | 0,77 | 0,90 |
| 70 | 0 | SR | meanT | Prevpara | 142,77 | 150,10 | 0,77 | 0,90 |
| 71 | 0 | meanR | meanT | Prevpara | 142,77 | 150,10 | 0,77 | 0,90 |
| 72 | LNlate | JR | ST | Prevpara | 142,78 | 151,57 | 0,80 | 0,91 |
| 73 | LNlate | JR | meanT | Prevpara | 142,78 | 151,57 | 0,80 | 0,91 |
| 74 | Ldens | SR | ST | Prevpara | 142,78 | 151,58 | 0,79 | 0,90 |
| 75 | Ldens | meanR | ST | Prevpara | 142,78 | 151,58 | 0,79 | 0,90 |
| 76 | Ldens | SR | meanT | Prevpara | 142,78 | 151,58 | 0,79 | 0,90 |
| 77 | Ldens | meanR | meanT | Prevpara | 142,78 | 151,58 | 0,79 | 0,90 |
| 78 | 0 | JR | ST | Prevpara | 142,79 | 150,12 | 0,77 | 0,91 |
| 79 | 0 | JR | meanT | Prevpara | 142,79 | 150,12 | 0,77 | 0,91 |
| 80 | LN | 0 | ST | Prevpara | 143,09 | 150,42 | 0,79 | 0,91 |
| 81 | LN | 0 | meanT | Prevpara | 143,09 | 150,42 | 0,79 | 0,91 |
| 82 | Ldens | AR | ST | Prevpara | 143,11 | 151,91 | 0,78 | 0,91 |
| 83 | Ldens | AR | meanT | Prevpara | 143,11 | 151,91 | 0,78 | 0,91 |
| 84 | LNlate | AR | ST | Prevpara | 143,25 | 152,05 | 0,79 | 0,91 |
| 85 | LNlate | AR | meanT | Prevpara | 143,25 | 152,05 | 0,79 | 0,91 |
| 86 | 0 | AR | ST | Prevpara | 143,37 | 150,70 | 0,77 | 0,91 |
| 87 | 0 | AR | meanT | Prevpara | 143,37 | 150,70 | 0,77 | 0,91 |
| 88 | Nymphs | 0 | ST | Prevpara | 143,40 | 150,73 | 0,76 | 0,91 |
| 89 | Nymphs | 0 | meanT | Prevpara | 143,40 | 150,73 | 0,76 | 0,91 |
| 90 | LN | JR | ST | Prevpara | 144,24 | 153,03 | 0,79 | 0,91 |
| 91 | LN | JR | meanT | Prevpara | 144,24 | 153,03 | 0,79 | 0,91 |
| 92 | LN | SR | ST | Prevpara | 144,42 | 153,22 | 0,79 | 0,90 |
| 93 | LN | meanR | ST | Prevpara | 144,42 | 153,22 | 0,79 | 0,90 |
| 94 | LN | SR | meanT | Prevpara | 144,42 | 153,22 | 0,79 | 0,90 |
| 95 | LN | meanR | meanT | Prevpara | 144,42 | 153,22 | 0,79 | 0,90 |
| 96 | Nymphs | JR | ST | Prevpara | 144,62 | 153,42 | 0,76 | 0,91 |
| 97 | Nymphs | JR | meanT | Prevpara | 144,62 | 153,42 | 0,76 | 0,91 |
| 98 | Nymphs | SR | ST | Prevpara | 144,65 | 153,44 | 0,77 | 0,91 |
| 99 | Nymphs | meanR | ST | Prevpara | 144,65 | 153,44 | 0,77 | 0,91 |
| 100 | Nymphs | SR | meanT | Prevpara | 144,65 | 153,44 | 0,77 | 0,91 |
| 101 | Nymphs | meanR | meanT | Prevpara | 144,65 | 153,44 | 0,77 | 0,91 |
| 102 | LN | AR | ST | Prevpara | 144,85 | 153,65 | 0,79 | 0,91 |
| 103 | LN | AR | meanT | Prevpara | 144,85 | 153,65 | 0,79 | 0,91 |
| 104 | Nymphs | AR | ST | Prevpara | 145,22 | 154,02 | 0,76 | 0,91 |
| 105 | Nymphs | AR | meanT | Prevpara | 145,22 | 154,02 | 0,76 | 0,91 |
| 106 | LNlate | 0 | ST | 0 | 153,23 | 159,10 | 0,66 | 0,91 |

| | | | | | | | | |
|---|---|---|---|---|---|---|---|---|
| 107 | LNlate | 0 | meanT | 0 | 153,23 | 159,10 | 0,66 | 0,91 |
| 108 | Ndens | JR | AT | Prevpara | 153,88 | 162,68 | 0,69 | 0,92 |
| 109 | Ldens | JR | AT | Prevpara | 153,89 | 162,68 | 0,69 | 0,91 |
| 110 | Larvae | JR | AT | Prevpara | 154,10 | 162,90 | 0,74 | 0,92 |
| 111 | Ldens | SR | JT | 0 | 154,32 | 161,64 | 0,58 | 0,90 |
| 112 | Ldens | meanR | JT | 0 | 154,32 | 161,64 | 0,58 | 0,90 |
| 113 | LNlate | 0 | JT | 0 | 154,34 | 160,20 | 0,67 | 0,91 |
| 114 | Larvae | SR | ST | 0 | 154,50 | 161,83 | 0,70 | 0,91 |
| 115 | Larvae | meanR | ST | 0 | 154,50 | 161,83 | 0,70 | 0,91 |
| 116 | Larvae | SR | meanT | 0 | 154,50 | 161,83 | 0,70 | 0,91 |
| 117 | Larvae | meanR | meanT | 0 | 154,50 | 161,83 | 0,70 | 0,91 |
| 118 | Ldens | SR | ST | 0 | 154,59 | 161,92 | 0,57 | 0,90 |
| 119 | Ldens | meanR | ST | 0 | 154,59 | 161,92 | 0,57 | 0,90 |
| 120 | Ldens | SR | meanT | 0 | 154,59 | 161,92 | 0,57 | 0,90 |
| 121 | Ldens | meanR | meanT | 0 | 154,59 | 161,92 | 0,57 | 0,90 |
| 122 | Larvae | SR | JT | 0 | 154,70 | 162,03 | 0,73 | 0,92 |
| 123 | Larvae | meanR | JT | 0 | 154,70 | 162,03 | 0,73 | 0,92 |
| 124 | LNlate | SR | ST | 0 | 154,83 | 162,16 | 0,67 | 0,92 |
| 125 | LNlate | meanR | ST | 0 | 154,83 | 162,16 | 0,67 | 0,92 |
| 126 | LNlate | SR | meanT | 0 | 154,83 | 162,16 | 0,67 | 0,92 |
| 127 | LNlate | meanR | meanT | 0 | 154,83 | 162,16 | 0,67 | 0,92 |
| 128 | Ldens | 0 | ST | 0 | 154,86 | 160,72 | 0,50 | 0,88 |
| 129 | Ldens | 0 | meanT | 0 | 154,86 | 160,72 | 0,50 | 0,88 |
| 130 | LNlate | AR | ST | 0 | 154,99 | 162,32 | 0,67 | 0,91 |
| 131 | LNlate | AR | meanT | 0 | 154,99 | 162,32 | 0,67 | 0,91 |
| 132 | LNlate | JR | ST | 0 | 155,21 | 162,54 | 0,66 | 0,91 |
| 133 | LNlate | JR | meanT | 0 | 155,21 | 162,54 | 0,66 | 0,91 |
| 134 | LNlate | SR | JT | 0 | 155,36 | 162,69 | 0,70 | 0,92 |
| 135 | LNlate | meanR | JT | 0 | 155,36 | 162,69 | 0,70 | 0,92 |
| 136 | 0 | JR | AT | Prevpara | 155,38 | 162,71 | 0,68 | 0,91 |
| 137 | Ldens | AR | ST | 0 | 155,51 | 162,84 | 0,54 | 0,89 |
| 138 | Ldens | AR | meanT | 0 | 155,51 | 162,84 | 0,54 | 0,89 |
| 139 | Ldens | 0 | JT | 0 | 155,65 | 161,51 | 0,49 | 0,88 |
| 140 | LNlate | JR | JT | 0 | 155,80 | 163,13 | 0,67 | 0,91 |
| 141 | LNlate | JR | AT | Prevpara | 155,80 | 164,59 | 0,70 | 0,91 |
| 142 | Nymphs | JR | AT | Prevpara | 156,14 | 164,93 | 0,64 | 0,92 |
| 143 | Larvae | 0 | ST | 0 | 156,28 | 162,14 | 0,64 | 0,89 |
| 144 | Larvae | 0 | meanT | 0 | 156,28 | 162,14 | 0,64 | 0,89 |
| 145 | LNlate | AR | JT | 0 | 156,32 | 163,65 | 0,67 | 0,91 |
| 146 | Ldens | AR | AT | Prevpara | 156,33 | 165,12 | 0,66 | 0,91 |
| 147 | Larvae | AR | ST | 0 | 156,57 | 163,90 | 0,67 | 0,89 |
| 148 | Larvae | AR | meanT | 0 | 156,57 | 163,90 | 0,67 | 0,89 |
| 149 | Ldens | JR | ST | 0 | 156,75 | 164,08 | 0,50 | 0,88 |
| 150 | Ldens | JR | meanT | 0 | 156,75 | 164,08 | 0,50 | 0,88 |
| 151 | Ndens | AR | AT | Prevpara | 156,96 | 165,76 | 0,66 | 0,93 |
| 152 | Ldens | AR | JT | 0 | 157,06 | 164,39 | 0,52 | 0,88 |
| 153 | LN | JR | AT | Prevpara | 157,17 | 165,97 | 0,70 | 0,91 |
| 154 | Ldens | AR | AT | 0 | 157,17 | 164,50 | 0,57 | 0,91 |
| 155 | Ldens | JR | JT | 0 | 157,55 | 164,88 | 0,49 | 0,88 |
| 156 | Larvae | AR | AT | Prevpara | 157,65 | 166,45 | 0,71 | 0,92 |
| 157 | Larvae | JR | ST | 0 | 158,22 | 165,55 | 0,64 | 0,89 |
| 158 | Larvae | JR | meanT | 0 | 158,22 | 165,55 | 0,64 | 0,89 |
| 159 | Ldens | JR | AT | 0 | 158,25 | 165,58 | 0,53 | 0,90 |
| 160 | Larvae | 0 | JT | 0 | 158,96 | 164,83 | 0,65 | 0,88 |
| 161 | Larvae | AR | AT | 0 | 159,01 | 166,34 | 0,66 | 0,92 |
| 162 | LNlate | JR | AT | 0 | 159,35 | 166,68 | 0,63 | 0,92 |
| 163 | 0 | AR | AT | Prevpara | 159,74 | 167,07 | 0,66 | 0,92 |
| 164 | LNlate | AR | AT | Prevpara | 159,82 | 168,62 | 0,69 | 0,92 |
| 165 | Ldens | 0 | AT | Prevpara | 160,02 | 167,35 | 0,62 | 0,91 |
| 166 | Larvae | JR | JT | 0 | 160,29 | 167,61 | 0,65 | 0,88 |
| 167 | Larvae | JR | AT | 0 | 160,31 | 167,63 | 0,62 | 0,91 |
| 168 | Larvae | AR | JT | 0 | 160,48 | 167,81 | 0,67 | 0,88 |
| 169 | LNlate | AR | AT | 0 | 160,49 | 167,82 | 0,65 | 0,92 |
| 170 | Nymphs | AR | AT | Prevpara | 161,11 | 169,90 | 0,64 | 0,93 |

| | | | | | | | | |
|---|---|---|---|---|---|---|---|---|
| 171 | Ldens | 0 | AT | 0 | 161,14 | 167,00 | 0,50 | 0,90 |
| 172 | LN | AR | AT | Prevpara | 161,25 | 170,05 | 0,69 | 0,92 |
| 173 | Ldens | SR | AT | Prevpara | 162,00 | 170,80 | 0,62 | 0,91 |
| 174 | Ldens | meanR | AT | Prevpara | 162,00 | 170,80 | 0,62 | 0,91 |
| 175 | Ldens | SR | AT | 0 | 162,30 | 169,63 | 0,53 | 0,91 |
| 176 | Ldens | meanR | AT | 0 | 162,30 | 169,63 | 0,53 | 0,91 |
| 177 | Ndens | AR | AT | 0 | 162,48 | 169,81 | 0,54 | 0,92 |
| 178 | LN | SR | ST | 0 | 162,58 | 169,91 | 0,66 | 0,90 |
| 179 | LN | meanR | ST | 0 | 162,58 | 169,91 | 0,66 | 0,90 |
| 180 | LN | SR | meanT | 0 | 162,58 | 169,91 | 0,66 | 0,90 |
| 181 | LN | meanR | meanT | 0 | 162,58 | 169,91 | 0,66 | 0,90 |
| 182 | LN | SR | JT | 0 | 162,93 | 170,26 | 0,70 | 0,90 |
| 183 | LN | meanR | JT | 0 | 162,93 | 170,26 | 0,70 | 0,90 |
| 184 | Ndens | SR | ST | 0 | 163,02 | 170,35 | 0,54 | 0,92 |
| 185 | Ndens | meanR | ST | 0 | 163,02 | 170,35 | 0,54 | 0,92 |
| 186 | Ndens | SR | meanT | 0 | 163,02 | 170,35 | 0,54 | 0,92 |
| 187 | Ndens | meanR | meanT | 0 | 163,02 | 170,35 | 0,54 | 0,92 |
| 188 | Ndens | 0 | AT | Prevpara | 163,19 | 170,51 | 0,61 | 0,92 |
| 189 | LN | AR | ST | 0 | 163,91 | 171,24 | 0,63 | 0,88 |
| 190 | LN | AR | meanT | 0 | 163,91 | 171,24 | 0,63 | 0,88 |
| 191 | LN | 0 | ST | 0 | 163,97 | 169,83 | 0,57 | 0,87 |
| 192 | LN | 0 | meanT | 0 | 163,97 | 169,83 | 0,57 | 0,87 |
| 193 | LNlate | 0 | AT | Prevpara | 164,07 | 171,40 | 0,66 | 0,92 |
| 194 | LNlate | 0 | AT | 0 | 164,44 | 170,30 | 0,61 | 0,92 |
| 195 | Ndens | JR | AT | 0 | 164,68 | 172,00 | 0,48 | 0,91 |
| 196 | 0 | AR | AT | 0 | 164,73 | 170,60 | 0,55 | 0,92 |
| 197 | LN | AR | AT | 0 | 164,79 | 172,12 | 0,63 | 0,91 |
| 198 | 0 | SR | ST | 0 | 165,05 | 170,91 | 0,54 | 0,91 |
| 199 | 0 | meanR | ST | 0 | 165,05 | 170,91 | 0,54 | 0,91 |
| 200 | 0 | SR | meanT | 0 | 165,05 | 170,91 | 0,54 | 0,91 |
| 201 | 0 | meanR | meanT | 0 | 165,05 | 170,91 | 0,54 | 0,91 |
| 202 | Ndens | SR | AT | Prevpara | 165,16 | 173,96 | 0,62 | 0,92 |
| 203 | Ndens | meanR | AT | Prevpara | 165,16 | 173,96 | 0,62 | 0,92 |
| 204 | Larvae | 0 | AT | Prevpara | 165,22 | 172,55 | 0,68 | 0,91 |
| 205 | LNlate | SR | AT | Prevpara | 165,55 | 174,34 | 0,67 | 0,91 |
| 206 | LNlate | meanR | AT | Prevpara | 165,55 | 174,34 | 0,67 | 0,91 |
| 207 | Ndens | SR | JT | 0 | 165,66 | 172,99 | 0,55 | 0,92 |
| 208 | Ndens | meanR | JT | 0 | 165,66 | 172,99 | 0,55 | 0,92 |
| 209 | Ndens | 0 | ST | 0 | 165,69 | 171,56 | 0,42 | 0,89 |
| 210 | Ndens | 0 | meanT | 0 | 165,69 | 171,56 | 0,42 | 0,89 |
| 211 | LN | JR | ST | 0 | 165,80 | 173,13 | 0,58 | 0,86 |
| 212 | LN | JR | meanT | 0 | 165,80 | 173,13 | 0,58 | 0,86 |
| 213 | 0 | JR | AT | 0 | 165,82 | 171,69 | 0,48 | 0,91 |
| 214 | LN | JR | AT | 0 | 166,14 | 173,47 | 0,56 | 0,90 |
| 215 | Nymphs | AR | AT | 0 | 166,20 | 173,52 | 0,53 | 0,92 |
| 216 | LN | 0 | JT | 0 | 166,20 | 172,06 | 0,60 | 0,85 |
| 217 | 0 | 0 | ST | 0 | 166,37 | 170,77 | 0,43 | 0,88 |
| 218 | 0 | 0 | meanT | 0 | 166,37 | 170,77 | 0,43 | 0,88 |
| 219 | 0 | 0 | AT | Prevpara | 166,43 | 172,29 | 0,63 | 0,91 |
| 220 | LNlate | SR | AT | 0 | 166,44 | 173,77 | 0,61 | 0,92 |
| 221 | LNlate | meanR | AT | 0 | 166,44 | 173,77 | 0,61 | 0,92 |
| 222 | Ndens | AR | ST | 0 | 166,46 | 173,79 | 0,47 | 0,89 |
| 223 | Ndens | AR | meanT | 0 | 166,46 | 173,79 | 0,47 | 0,89 |
| 224 | 0 | AR | ST | 0 | 166,93 | 172,79 | 0,48 | 0,89 |
| 225 | 0 | AR | meanT | 0 | 166,93 | 172,79 | 0,48 | 0,89 |
| 226 | Nymphs | SR | ST | 0 | 166,98 | 174,30 | 0,55 | 0,91 |
| 227 | Nymphs | meanR | ST | 0 | 166,98 | 174,30 | 0,55 | 0,91 |
| 228 | Nymphs | SR | meanT | 0 | 166,98 | 174,30 | 0,55 | 0,91 |
| 229 | Nymphs | meanR | meanT | 0 | 166,98 | 174,30 | 0,55 | 0,91 |
| 230 | Nymphs | JR | AT | 0 | 167,17 | 174,50 | 0,45 | 0,91 |
| 231 | Larvae | SR | AT | Prevpara | 167,17 | 175,96 | 0,68 | 0,91 |
| 232 | Larvae | meanR | AT | Prevpara | 167,17 | 175,96 | 0,68 | 0,91 |
| 233 | LN | AR | JT | 0 | 167,29 | 174,62 | 0,64 | 0,86 |
| 234 | 0 | SR | JT | 0 | 167,42 | 173,29 | 0,56 | 0,91 |

| | | | | | | | | |
|---|---|---|---|---|---|---|---|---|
| 235 | 0 | meanR | JT | 0 | 167,42 | 173,29 | 0,56 | 0,91 |
| 236 | Ndens | JR | ST | 0 | 167,68 | 175,01 | 0,42 | 0,89 |
| 237 | Ndens | JR | meanT | 0 | 167,68 | 175,01 | 0,42 | 0,89 |
| 238 | Ldens | SR | 0 | 0 | 167,72 | 173,58 | 0,45 | 0,90 |
| 239 | Ldens | meanR | 0 | 0 | 167,72 | 173,58 | 0,45 | 0,90 |
| 240 | Larvae | 0 | AT | 0 | 167,89 | 173,75 | 0,58 | 0,91 |
| 241 | LN | JR | JT | 0 | 167,98 | 175,31 | 0,60 | 0,85 |
| 242 | LN | 0 | AT | Prevpara | 168,12 | 175,45 | 0,65 | 0,91 |
| 243 | Nymphs | 0 | ST | 0 | 168,18 | 174,05 | 0,45 | 0,88 |
| 244 | Nymphs | 0 | meanT | 0 | 168,18 | 174,05 | 0,45 | 0,88 |
| 245 | 0 | SR | AT | Prevpara | 168,22 | 175,55 | 0,63 | 0,91 |
| 246 | 0 | meanR | AT | Prevpara | 168,22 | 175,55 | 0,63 | 0,91 |
| 247 | Ldens | AR | 0 | 0 | 168,32 | 174,19 | 0,44 | 0,89 |
| 248 | 0 | JR | ST | 0 | 168,34 | 174,21 | 0,43 | 0,88 |
| 249 | 0 | JR | meanT | 0 | 168,34 | 174,21 | 0,43 | 0,88 |
| 250 | Nymphs | 0 | AT | Prevpara | 168,40 | 175,73 | 0,62 | 0,91 |
| 251 | Ldens | 0 | 0 | 0 | 168,68 | 173,07 | 0,42 | 0,89 |
| 252 | Larvae | SR | AT | 0 | 168,73 | 176,06 | 0,61 | 0,92 |
| 253 | Larvae | meanR | AT | 0 | 168,73 | 176,06 | 0,61 | 0,92 |
| 254 | Nymphs | AR | ST | 0 | 168,79 | 176,12 | 0,49 | 0,89 |
| 255 | Nymphs | AR | meanT | 0 | 168,79 | 176,12 | 0,49 | 0,89 |
| 256 | Nymphs | SR | JT | 0 | 168,89 | 176,22 | 0,58 | 0,90 |
| 257 | Nymphs | meanR | JT | 0 | 168,89 | 176,22 | 0,58 | 0,90 |
| 258 | Ldens | AR | 0 | Prevpara | 169,54 | 176,87 | 0,48 | 0,89 |
| 259 | Ldens | 0 | 0 | Prevpara | 169,54 | 175,41 | 0,48 | 0,89 |
| 260 | Ldens | SR | 0 | Prevpara | 169,65 | 176,98 | 0,46 | 0,90 |
| 261 | Ldens | meanR | 0 | Prevpara | 169,65 | 176,98 | 0,46 | 0,90 |
| 262 | Ndens | SR | AT | 0 | 169,79 | 177,12 | 0,48 | 0,93 |
| 263 | Ndens | meanR | AT | 0 | 169,79 | 177,12 | 0,48 | 0,93 |
| 264 | Ndens | 0 | AT | 0 | 169,86 | 175,72 | 0,43 | 0,91 |
| 265 | LN | SR | AT | Prevpara | 169,97 | 178,76 | 0,66 | 0,91 |
| 266 | LN | meanR | AT | Prevpara | 169,97 | 178,76 | 0,66 | 0,91 |
| 267 | Nymphs | JR | ST | 0 | 170,16 | 177,49 | 0,45 | 0,88 |
| 268 | Nymphs | JR | meanT | 0 | 170,16 | 177,49 | 0,45 | 0,88 |
| 269 | Nymphs | SR | AT | Prevpara | 170,18 | 178,98 | 0,62 | 0,91 |
| 270 | Nymphs | meanR | AT | Prevpara | 170,18 | 178,98 | 0,62 | 0,91 |
| 271 | Ldens | JR | 0 | 0 | 170,33 | 176,20 | 0,42 | 0,89 |
| 272 | Ndens | 0 | JT | 0 | 170,70 | 176,56 | 0,42 | 0,88 |
| 273 | 0 | 0 | JT | 0 | 170,80 | 175,20 | 0,42 | 0,87 |
| 274 | Ldens | JR | 0 | Prevpara | 170,95 | 178,28 | 0,49 | 0,89 |
| 275 | Ndens | JR | JT | 0 | 171,66 | 178,98 | 0,41 | 0,88 |
| 276 | Nymphs | 0 | JT | 0 | 171,77 | 177,63 | 0,46 | 0,87 |
| 277 | 0 | JR | JT | 0 | 172,22 | 178,08 | 0,42 | 0,88 |
| 278 | 0 | AR | JT | 0 | 172,31 | 178,17 | 0,45 | 0,88 |
| 279 | Ndens | AR | JT | 0 | 172,31 | 179,64 | 0,45 | 0,88 |
| 280 | 0 | 0 | AT | 0 | 172,86 | 177,26 | 0,45 | 0,91 |
| 281 | Nymphs | JR | JT | 0 | 173,23 | 180,56 | 0,46 | 0,87 |
| 282 | LN | 0 | AT | 0 | 173,33 | 179,19 | 0,53 | 0,90 |
| 283 | Nymphs | AR | JT | 0 | 173,34 | 180,67 | 0,49 | 0,88 |
| 284 | 0 | SR | AT | 0 | 173,62 | 179,48 | 0,50 | 0,92 |
| 285 | 0 | meanR | AT | 0 | 173,62 | 179,48 | 0,50 | 0,92 |
| 286 | LN | SR | AT | 0 | 174,01 | 181,34 | 0,57 | 0,91 |
| 287 | LN | meanR | AT | 0 | 174,01 | 181,34 | 0,57 | 0,91 |
| 288 | Nymphs | 0 | AT | 0 | 174,84 | 180,70 | 0,44 | 0,91 |
| 289 | Nymphs | SR | AT | 0 | 175,60 | 182,93 | 0,49 | 0,92 |
| 290 | Nymphs | meanR | AT | 0 | 175,60 | 182,93 | 0,49 | 0,92 |
| 291 | LNlate | SR | 0 | 0 | 184,88 | 190,75 | 0,43 | 0,88 |
| 292 | LNlate | meanR | 0 | 0 | 184,88 | 190,75 | 0,43 | 0,88 |
| 293 | LNlate | AR | 0 | 0 | 185,05 | 190,91 | 0,42 | 0,87 |
| 294 | LNlate | 0 | 0 | 0 | 185,13 | 189,53 | 0,38 | 0,87 |
| 295 | LNlate | AR | 0 | Prevpara | 186,38 | 193,71 | 0,42 | 0,86 |
| 296 | LNlate | 0 | 0 | Prevpara | 186,41 | 192,27 | 0,39 | 0,86 |
| 297 | LNlate | JR | 0 | 0 | 186,72 | 192,58 | 0,39 | 0,87 |
| 298 | LNlate | SR | 0 | Prevpara | 186,79 | 194,11 | 0,42 | 0,88 |

| | | | | | | | | |
|---|---|---|---|---|---|---|---|---|
| 299 | LNlate | meanR | 0 | Prevpara | 186,79 | 194,11 | 0,42 | 0,88 |
| 300 | LNlate | JR | 0 | Prevpara | 187,74 | 195,07 | 0,40 | 0,85 |
| 301 | Larvae | SR | 0 | 0 | 188,42 | 194,28 | 0,39 | 0,88 |
| 302 | Larvae | meanR | 0 | 0 | 188,42 | 194,28 | 0,39 | 0,88 |
| 303 | Larvae | SR | 0 | Prevpara | 189,96 | 197,29 | 0,39 | 0,87 |
| 304 | Larvae | meanR | 0 | Prevpara | 189,96 | 197,29 | 0,39 | 0,87 |
| 305 | Larvae | AR | 0 | Prevpara | 190,17 | 197,50 | 0,40 | 0,84 |
| 306 | Ndens | SR | 0 | Prevpara | 190,22 | 197,55 | 0,33 | 0,88 |
| 307 | Ndens | meanR | 0 | Prevpara | 190,22 | 197,55 | 0,33 | 0,88 |
| 308 | Ndens | SR | 0 | 0 | 190,27 | 196,13 | 0,32 | 0,90 |
| 309 | Ndens | meanR | 0 | 0 | 190,27 | 196,13 | 0,32 | 0,90 |
| 310 | LN | AR | 0 | Prevpara | 190,30 | 197,62 | 0,46 | 0,81 |
| 311 | LN | SR | 0 | 0 | 190,69 | 196,56 | 0,41 | 0,85 |
| 312 | LN | meanR | 0 | 0 | 190,69 | 196,56 | 0,41 | 0,85 |
| 313 | LN | SR | 0 | Prevpara | 191,52 | 198,85 | 0,41 | 0,83 |
| 314 | LN | meanR | 0 | Prevpara | 191,52 | 198,85 | 0,41 | 0,83 |
| 315 | Nymphs | AR | 0 | Prevpara | 191,66 | 198,99 | 0,39 | 0,80 |
| 316 | Ndens | AR | 0 | Prevpara | 191,78 | 199,11 | 0,32 | 0,84 |
| 317 | Larvae | AR | 0 | 0 | 191,87 | 197,73 | 0,31 | 0,84 |
| 318 | Larvae | 0 | 0 | Prevpara | 192,27 | 198,13 | 0,39 | 0,83 |
| 319 | 0 | AR | 0 | Prevpara | 192,31 | 198,17 | 0,27 | 0,82 |
| 320 | LN | AR | 0 | 0 | 192,40 | 198,27 | 0,38 | 0,81 |
| 321 | Nymphs | SR | 0 | Prevpara | 192,57 | 199,90 | 0,38 | 0,82 |
| 322 | Nymphs | meanR | 0 | Prevpara | 192,57 | 199,90 | 0,38 | 0,82 |
| 323 | LN | 0 | 0 | Prevpara | 192,90 | 198,76 | 0,42 | 0,80 |
| 324 | 0 | SR | 0 | Prevpara | 193,00 | 198,86 | 0,27 | 0,85 |
| 325 | 0 | meanR | 0 | Prevpara | 193,00 | 198,86 | 0,27 | 0,85 |
| 326 | Ndens | 0 | 0 | Prevpara | 193,08 | 198,94 | 0,29 | 0,84 |
| 327 | Larvae | JR | 0 | Prevpara | 193,09 | 200,42 | 0,40 | 0,83 |
| 328 | Nymphs | 0 | 0 | Prevpara | 193,16 | 199,02 | 0,35 | 0,77 |
| 329 | LN | JR | 0 | Prevpara | 193,19 | 200,52 | 0,46 | 0,80 |
| 330 | Nymphs | SR | 0 | 0 | 193,47 | 199,34 | 0,36 | 0,86 |
| 331 | Nymphs | meanR | 0 | 0 | 193,47 | 199,34 | 0,36 | 0,86 |
| 332 | 0 | SR | 0 | 0 | 193,58 | 197,98 | 0,27 | 0,88 |
| 333 | 0 | meanR | 0 | 0 | 193,58 | 197,98 | 0,27 | 0,88 |
| 334 | Nymphs | JR | 0 | Prevpara | 193,88 | 201,21 | 0,37 | 0,77 |
| 335 | 0 | 0 | 0 | Prevpara | 194,28 | 198,68 | 0,21 | 0,80 |
| 336 | Ndens | JR | 0 | Prevpara | 194,42 | 201,75 | 0,31 | 0,83 |
| 337 | 0 | JR | 0 | Prevpara | 194,94 | 200,80 | 0,24 | 0,80 |
| 338 | Larvae | 0 | 0 | 0 | 195,16 | 199,55 | 0,23 | 0,83 |
| 339 | Nymphs | AR | 0 | 0 | 196,29 | 202,16 | 0,28 | 0,83 |
| 340 | LN | 0 | 0 | 0 | 196,69 | 201,09 | 0,24 | 0,78 |
| 341 | Larvae | JR | 0 | 0 | 196,75 | 202,61 | 0,23 | 0,82 |
| 342 | Ndens | AR | 0 | 0 | 196,93 | 202,80 | 0,19 | 0,85 |
| 343 | 0 | AR | 0 | 0 | 196,99 | 201,38 | 0,17 | 0,84 |
| 344 | LN | JR | 0 | 0 | 197,82 | 203,68 | 0,26 | 0,78 |
| 345 | Nymphs | 0 | 0 | 0 | 199,38 | 203,78 | 0,11 | 0,79 |
| 346 | Ndens | 0 | 0 | 0 | 199,68 | 204,08 | 0,07 | 0,84 |
| 347 | 0 | 0 | 0 | 0 | 200,56 | 203,50 | 0,00 | 0,82 |
| 348 | Nymphs | JR | 0 | 0 | 200,93 | 206,79 | 0,13 | 0,80 |
| 349 | Ndens | JR | 0 | 0 | 201,59 | 207,46 | 0,07 | 0,84 |
| 350 | 0 | JR | 0 | 0 | 202,10 | 206,50 | 0,01 | 0,82 |
