## Supplementary material for "Temporal variation and drivers of *Ixodiphagus hookeri* parasitisation in *Ixodes ricinus* ticks in northern Europe": Table S2

**Table S2.** Study site and year specific *Ixodiphagus hookeri* parasitisation prevalence in questing *Ixodes ricinus* nymphs on Seili Island.

| Year | Study site | Parasitized ticks | Analysed samples | Parasitization percentage | 95 % confidence interval | |
|---|---|---|---|---|---|---|
| | | | | | Lower | Upper |
| 2014 | A1 | 0 | 13 | 0 | 0 | 0 |
| 2015 | A1 | 0 | 3 | 0 | 0 | 0 |
| 2016 | A1 | 0 | 21 | 0 | 0 | 0 |
| 2017 | A1 | 6 | 41 | 14,6 | 6,9 | 28,4 |
| 2018 | A1 | 2 | 47 | 4,3 | 1,1 | 14,2 |
| 2019 | A1 | 19 | 90 | 21,1 | 9,5 | 30,6 |
| 2020 | A1 | 7 | 59 | 11,9 | 5,9 | 22,5 |
| 2021 | A1 | 7 | 64 | 10,9 | 5,4 | 20,9 |
| 2014 | C2 | 0 | 94 | 0 | 0 | 0 |
| 2015 | C2 | 0 | 13 | 0 | 0 | 0 |
| 2016 | C2 | 0 | 61 | 0 | 0 | 0 |
| 2017 | C2 | 0 | 7 | 0 | 0 | 0 |
| 2018 | C2 | 3 | 38 | 7,9 | 2,7 | 20,8 |
| 2019 | C2 | 9 | 86 | 10,5 | 5,6 | 18,7 |
| 2020 | C2 | 5 | 162 | 2,7 | 1,3 | 7,0 |
| 2021 | C2 | 6 | 171 | 3,5 | 1,6 | 7,4 |
| 2014 | C3 | 1 | 56 | 1,8 | 0,3 | 9,4 |
| 2015 | C3 | 2 | 17 | 11,8 | 3,3 | 34,3 |
| 2016 | C3 | 3 | 54 | 5,6 | 1,9 | 15,1 |
| 2017 | C3 | 3 | 38 | 7,9 | 2,7 | 20,8 |
| 2018 | C3 | 4 | 42 | 9,5 | 3,8 | 22,1 |
| 2019 | C3 | 25 | 66 | 37,9 | 27,1 | 49,9 |
| 2020 | C3 | 12 | 72 | 16,7 | 9,8 | 27,0 |
| 2021 | C3 | 3 | 52 | 5,8 | 2,0 | 15,6 |
| 2014 | D1 | 1 | 74 | 1,4 | 0,2 | 7,3 |
| 2015 | D1 | 1 | 20 | 5 | 0,9 | 23,6 |
| 2016 | D1 | 5 | 106 | 4,7 | 2,0 | 10,6 |
| 2017 | D1 | 3 | 79 | 3,8 | 1,3 | 10,6 |
| 2018 | D1 | 4 | 50 | 8,0 | 3,2 | 18,8 |
| 2019 | D1 | 18 | 68 | 26,5 | 17,5 | 38,0 |
| 2020 | D1 | 13 | 74 | 17,6 | 10,6 | 27,7 |
| 2021 | D1 | 12 | 99 | 3,5 | 7,0 | 20,0 |
